# Predictive control of electrophysiological network architecture using direct, single-node neurostimulation in humans

**DOI:** 10.1101/292748

**Authors:** Ankit N. Khambhati, Ari E. Kahn, Julia Costantini, Youssef Ezzyat, Ethan A. Solomon, Robert E. Gross, Barbara C. Jobst, Sameer A. Sheth, Kareem A. Zaghloul, Gregory Worrell, Sarah Seger, Bradley C. Lega, Shennan Weiss, Michael R. Sperling, Richard Gorniak, Sandhitsu R. Das, Joel M. Stein, Daniel S. Rizzuto, Michael J. Kahana, Timothy H. Lucas, Kathryn A. Davis, Joseph I. Tracy, Danielle S. Bassett

## Abstract

Chronically implantable neurostimulation devices are becoming a clinically viable option for treating patients with neurological disease and psychiatric disorders. Neurostimulation offers the ability to probe and manipulate distributed networks of interacting brain areas in dysfunctional circuits. Here, we use tools from network control theory to examine the dynamic reconfiguration of functionally interacting neuronal ensembles during targeted neurostimulation of cortical and subcortical brain structures. By integrating multi-modal intracranial recordings and diffusion tensor imaging from patients with drug-resistant epilepsy, we test hypothesized structural and functional rules that predict altered patterns of synchronized local field potentials. We demonstrate the ability to predictably reconfigure functional interactions depending on stimulation strength and location. Stimulation of areas with structurally weak connections largely modulates the functional hubness of downstream areas and concurrently propels the brain towards more difficult-to-reach dynamical states. By using focal perturbations to bridge large-scale structure, function, and markers of behavior, our findings suggest that stimulation may be tuned to influence different scales of network interactions driving cognition.

## 1. Introduction

Novel neurotechnologies capable of perturbing the physiological state of neural systems are rapidly gaining popularity for their potential to treat neurological disease and psychiatric disorders [87]. Chronically implantable devices that stimulate the human brain are clinically approved to treat Parkinson’s disease, essential tremor, dystonia, epilepsy, and obsessive-compulsive disorder and have been investigated for major depressive disorder and Tourette syndrome [66]. Recent human studies have investigated the ability for direct stimulation of cortical and subcortical structures to modulate biomarkers of memory [35, 47], visual perception [80, 97], language production [31], somatosensory perception [73], sensorimotor function [96], and subjective experience [37]. While neurostimulation is a promising interventional approach to modulate brain state, current practices of calibrating *where*, *when*, and *how* to stimulate the brain are “open-loop” and limited in efficacy – relying on manual and periodic tuning of device parameters to optimize therapy [71]. Automated, “closed-loop” approaches would augment the capability of current stimulation devices to dynamically adjust parameters based on the physiological state of the brain network, monitored in real-time [88]. Undoubtedly, the translational prospect of neurostimulation to manipulate brain networks that generate abnormal rhythms, dysrhythmias, or bursts of activity associated with dysfunction is promising. However, critical gaps in knowledge hinder the development of a robust control policy for next-generation implantable devices.

How does the architecture of the neural system mediate the effect of neurostimulation on neurophysiology and behavior? Network control theory [77] provides a mathematical framework for mapping the influence of a control signal on the dynamics of an interconnected system. When combined with graph modelling tools from network neuroscience [10], where *nodes* represent discrete brain regions and *edges* represent the structural connections between brain regions, control theoretic approaches can elucidate how the brain’s structural architecture of white matter fiber pathways shapes its ability to navigate through a repertoire of dynamical states [42]. Theoretical rules of *controllability* prescribe the trajectories through state space elicited by a given control signal [42, 15, 41], and begin to explain why one brain network may be more or less influential on brain dynamics than another [59]. Recent efforts to test control theoretic predictions of the relationship between controllability and brain activity have relied on *in silico* models in which neuronal ensembles are interlinked by structural connections measured by human neuroimaging [72]. Despite the promising convergence between theory and model simulation, empirical stimulation data bridging network control and neurophysiology are lacking.

Network control theory accounts for the structural connections that convey modulated brain activity to downstream regions in the network; however, it does not account for the functional rules that govern whether communication between brain regions can occur at a specific point in time. At the millimeter-scale, synchronous oscillations in the local field potential are thought to actively gate the transfer of information across the network [29, 27, 82, 38, 18] and are commonly observed during higher-order cognitive processing [26]. A functional (rather than structural) network representation of the coherence between different ensembles of neurons may capture dynamical states of communication [46, 33]. The neurophysiologic interpretation of these states can depend on the measured frequency range of the functional network [8, 85], which in turn implicates certain types of cells interacting over specific spatial scales [60]. Prior studies have examined how these functional networks may reconfigure during higher-order cognitive functions such as learning new skills [11, 13, 12, 69], forming memories [20], attending to the environment [84], and processing language [30]. Complimentary work also posits that reconfiguration of functional networks may underlie neurophysiological abnormalities in patients with epilepsy [58, 56], schizophrenia [9, 19], Parkinson’s disease [81, 75], and stroke [95, 40]. While these studies explain changes in functional network reconfiguration when the brain is perturbed *en masse*, a rigorously quantified map of functional network reconfiguration due to controlled, focal perturbation has not been attained.

Here we seek to elucidate the network control principles by which neurostimulation can alter function and behavior based on constraints prescribed by structural connectivity and spontaneous functional interactions. We measure the electrocorticogram (ECoG) in 94 drug-resistant epilepsy patients undergoing neurostimulation (**Fig. 1a-b**), and we construct structural networks from diffusion imaging data acquired in the same individuals. We also construct functional networks before and after individual stimulation trials using multitaper coherence between sensors [79] in distinct frequency bands [63, 57] (**Fig. 1c**), and we define brain state before and after stimulation using a previously validated biomarker of memory [35]. We test four hypotheses. First, we hypothesize that the strength and location of stimulation can differentially drive two separate modes of global versus local control over functional architecture [72]. Intuitively, stimulation to functional hubs – nodes that tend to interact strongly with the rest of the network – may have swiftly attenuated effects due to signal dispersion across many downstream regions, while stimulation to non-hubs may have more localized and targeted effects. Second, we hypothesize that regions with strong baseline functional interaction with the stimulation site are more likely to exhibit altered hub properties following stimulation than brain regions with weak functional interaction with the stimulation site, indicating a functional conduit of stimulation. Third, based on prior data [16], we hypothesize that these functional interactions – particularly in high frequency bands – co-localize with structural white matter networks (**Fig. 1d**). Fourth, we hypothesize that neurostimulation directed towards modal control points [42, 77], which tend to be structural non-hubs of a patient’s white matter network thereby minimizing signal dispersion, facilitate a stronger shift in dynamical state associated with memory encoding, a function that is altered in patients with epilepsy [45, 94, 1]. Collectively, these analyses will supply a roadmap of the impact of neurostimulation on network physiology, mediated by network structure, and provide fundamental mechanistic insight into the influence of neurostimulation on behavioral state.

**Figure 1:**
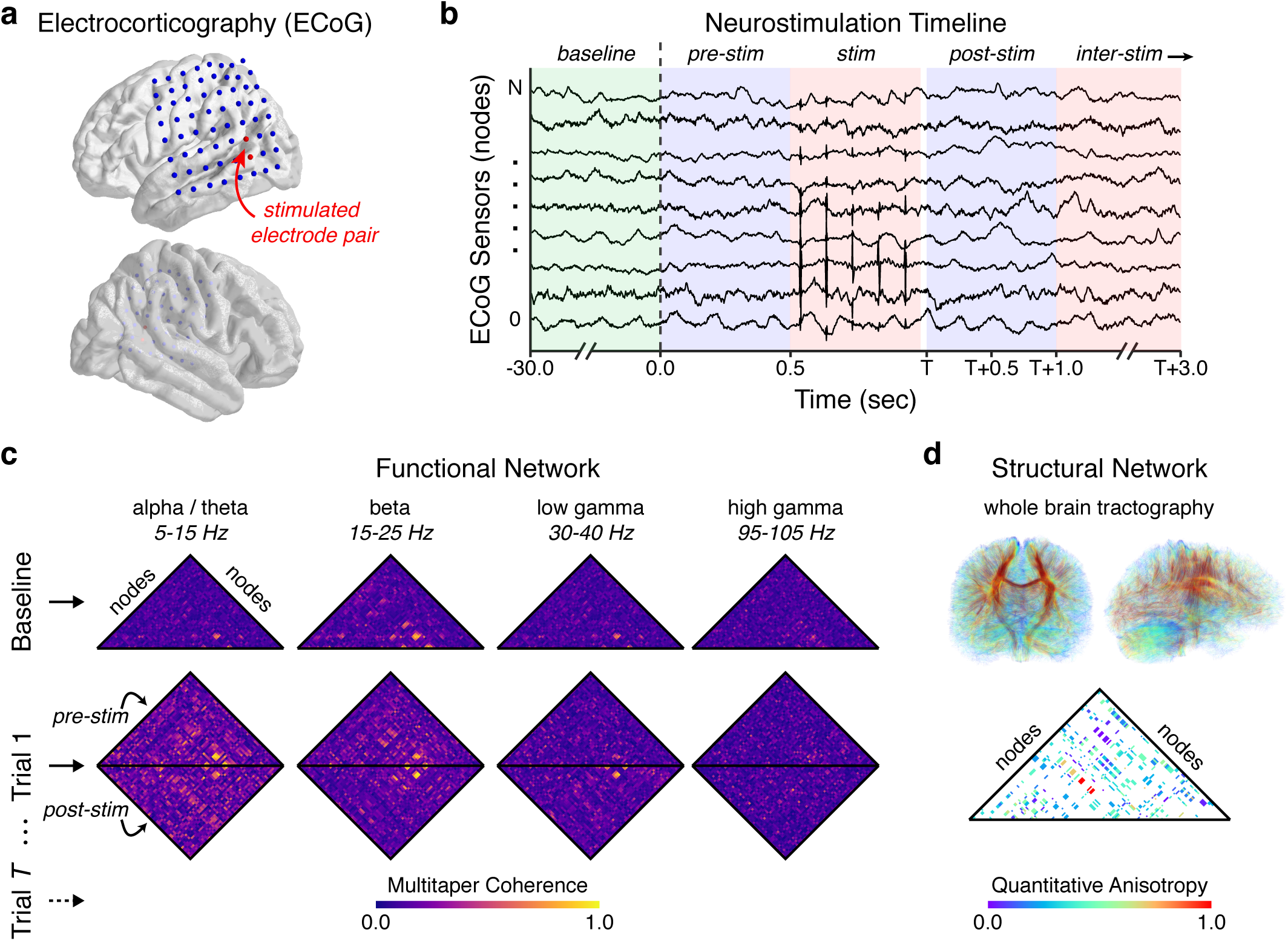
Measuring network response to targeted, intracranial neurostimulation. (**a**) We record the electrocorticogram (ECoG) in 94 patients with drugresistant epilepsy across 8 clinical institutions using intracranial sensors implanted in cortical and subcortical brain structures. To evoke a network response, we stimulate adjacent electrode pairs using a charge-balanced, biphasic current source with a square waveform of variable amplitude, frequency, and duration. (**b**) For each experimental session, we select a stimulation location and collect the following epochs of ECoG activity: (i) thirty seconds of baseline activity before any stimulation is given, (ii) one half-second of activity before a stimulation trial, and (iii) two consecutive and non-overlapping half-second windows of activity after a stimulation trial. A stimulation trial is defined by a combination of pulse frequency, amplitude, and duration, and consecutive stimulation trials are separated by an inter-stimulation interval drawn from a uniform random distribution ranging from 2.75 seconds to 3.25 seconds. (**c**) We measure the impact of neurostimulation on functional network architecture by constructing dynamic graph models in which intracranial sensors are represented by *nodes* and the functional interactions between intracranial sensors are represented by *edges*. To infer functional interactions, we calculate the multitaper coherence between each pair of ECoG signals in non-overlapping, half-second time windows for each baseline epoch, pre-stimulation epoch, and post-stimulation epoch in the following four frequency bands: (i) alpha/theta (5-15 Hz), (ii) beta (15-25 Hz), (iii) low gamma (30-40 Hz), and (iv) high gamma (95-105 Hz) [63, 57]. (**d**) To examine how structural connectivity constrains functional network reconfiguration to neurostimulation, we also construct a static graph model of the brain’s structural network by applying deterministic tractography to each subject’s diffusion tensor imaging data.

## 2. Results

### 2.1. Neurostimulation drives localized and distributed functional network reconfiguration

We first ask the question, “How does neurostimulation alter the architecture of functional brain networks?” Based on recent theoretical insights on the costs of forming and breaking connections in structural and functional brain networks [17, 21, 2], we expect stimulation to heterogeneously affect existing coherent interactions, strengthening some and weakening others. To test these expectations, we study three measures of network reconfiguration: two at the topological scale of nodes and one at the topological scale of edges (**Fig. 2a**). At the node scale, we first compute the strength, or average coherence, for each network node during the pre-stim epoch and post-stim epoch for each of the four coherence frequency bands. We next examine the *change in the mean of node strengths* and the *change in the variance of node strengths* between the pre-stim epoch and the post-stim epoch. Intuitively, a change in the mean of node strengths quantifies the likelihood that nodes exhibit greater frequency-specific functional interaction following stimulation, and a change in the variance of node strengths quantifies the likelihood that nodes exhibit greater heterogeneity in their degree of functional interaction with other nodes in the network. At the edge scale, we compute the *configuration similarity* [58]: a Pearson correlation between the vector of coherence weights during the pre-stim epoch and the vector of coherence weights during the post-stim epoch. Similarity values near 0 imply a greater change in the configuration of network coherences, and values near 1 imply a lesser change in the configuration of network coherences. We confirmed that the topological measurements at the node scale capture different reconfiguration phenomenon than the topological measurements at the edge scale by observing weak relationships between changes in the mean and variance of node strengths to configuration similarity (**Fig. S1**).

**Figure 2:**
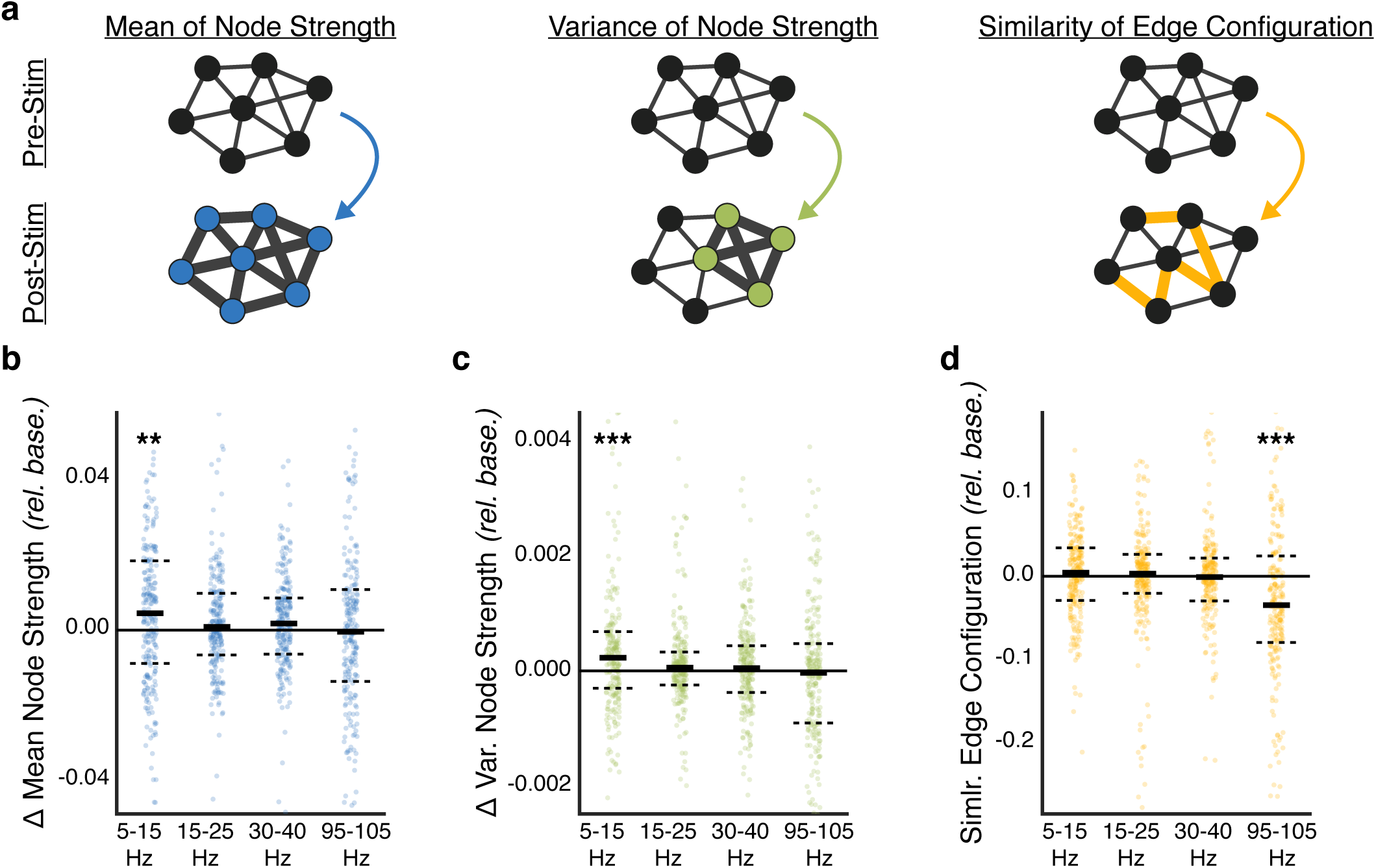
Control of frequency-specific functional network topology. (**a**) Does stimulation induce network reconfiguration at the scale of network nodes or at the scale of network edges? Shown here are different forms of network reconfiguration: two forms at the node scale and one form at the edge scale. At the node scale, stimulation may increase or decrease the overall functional interactions of a node with other nodes in the network, resulting in a change in the mean of node strengths and/or a change in the variance, or heterogeneity, of node strengths in the network. At the edge scale, stimulation may alter the configurational pattern of functional interactions underlying functional network topology. We measure edge scale change by computing a configuration similarity metric [58] of the pattern of network coherences between the pre-stim trial and the post-stim trial; values near 1 (or 0) imply a lesser (or greater) change in network configuration. (**b**) Difference in the change in mean of node strengths between stim epochs and baseline epochs. Change in the mean of node strengths is significantly greater during stimulation epochs than baseline epochs in the alpha/theta band (*p* < 0.01, corrected). (**c**) Difference in the change in variance of node strengths between stim epochs and baseline epochs. Change in the variance of node strengths is significantly greater during stimulation epochs than baseline epochs in the alpha/theta band (*p* < 0.001, corrected). Stimulation alters low frequency organization of the functional network at the scale of network nodes. (**d**) Difference in the configuration similarity of network edges between stim epochs and baseline epochs. Reconfiguration of functional interactions is significantly greater during stimulation epochs than baseline epochs in the high gamma band (*p* < 0.001, corrected). Stimulation alters high frequency organization of the functional network at the scale of network edges. Each observation is the average across epochs within a stimulation session of a single subject. Solid lines represent the median, and dashed lines represent the first and third quartiles. \**p* < 0.05, \*\**p* < 0.01, \*\*\**p* < 0.001.

Next, we test our expectation that stimulation heterogeneously affects existing coherent interactions, strengthening some and weakening others. We first study changes in the mean and variance of node strengths between the pre-stim epoch and post-stim epoch for each of the four coherence frequency bands (**Fig. 2b,c**). Using a Wilcoxon rank-sum test and Bonferroni correction for multiple comparisons, we examine whether node-level changes in the network 100ms after stimulation offset are any greater than passive changes observed over an equal duration of spontaneous activity at baseline, before any stimulation, across stimulation sessions over subjects. We find that stimulation leads to a significantly greater change in the mean of node strengths than expected at baseline in the alpha/theta band (*Z*(247) = 11448, *p* = 0.001) and to a non-significant change in the beta band (*Z*(247) = 14006, *p* = 0.68), in the low gamma band (*Z*(247) = 13131, *p* = 0.13), and in the high gamma band (*Z*(247) = 14883, *p* = 1.0). We also find that stimulation leads to a significantly greater change in the variance of node strengths than expected at baseline in the alpha/theta band (*Z*(247) = 10660, *p* = 0.0001) and to a non-significant change in the beta band (*Z*(247) = 13422, *p* = 0.24), in the low gamma band (*Z*(247) = 14137, *p* = 0.84), and in the high gamma band (*Z*(247) = 14086, *p* = 0.76). We find that these effects indeed persist and possibly strengthen in the beta band and low gamma band at least 600ms after stimulation offset (**Fig. S2**b,c). These results demonstrate that stimulation amenably alters functional network organization in lower alpha/theta band frequencies (5– 15 Hz) at the node scale. Specifically, we observe that nodes generally exhibit an increase in low frequency interaction following neurostimulation. However, changes in node strengths are also heterogeneously distributed across nodes in the network.

We next ask whether stimulation may still alter functional network topology at the edge scale. Using a Wilcoxon ranksum test and Bonferroni correction for multiple comparisons, we examine whether configurational changes in the network edges 100ms after stimulation offset are any greater than the passive change observed over an equal duration of spontaneous activity at baseline, before any stimulation, across stimulation sessions over subjects. We find that stimulation leads to a significantly lower configuration similarity (greater reconfiguration) than expected at baseline in the high gamma band (*Z*(247) = 9252, *p* = 2.3×10^−6^) and to a non-significant change in the alpha/theta band (*Z*(247) = 13543, *p* = 1.0), in the beta band (*Z*(247) = 12820, *p* = 1.0), and in the low gamma band (*Z*(247) = 13502, *p* = 1.0). We find that these effects indeed persist at least 600 ms after stimulation offset (**Fig. S2**d). These results demonstrate that stimulation amenably alters functional network organization in high gamma band frequencies (95–105 Hz) at the edge scale. Specifically, we observe that functional interactions undergo a change in their configurational pattern in high frequencies following neurostimulation.

### 2.2. Input energy differentially modulates topological scale of functional network response

Building on our observations of a complex, frequencydependent network response to stimulation, we next ask, “Do properties of the stimulation signal, such as amplitude, pulse frequency, and duration, influence the extent of functional network reconfiguration?” To answer this question, we iteratively cycle through stimulation parameters for each consecutive trial (**Fig. 3a**), and we compute the stimulation energy as the product between the three parameters (**Fig. 3b**). Based on prior observations of a relationship between stimulation energy and volume of tissue activated [25], we hypothesize that stronger stimulation input into the functional network will lead to more widespread change in functional architecture than weaker stimulation input, presumably by penetrating the network along short axonal fibers in the gray matter and long myelinated fibers in the white matter.

**Figure 3:**
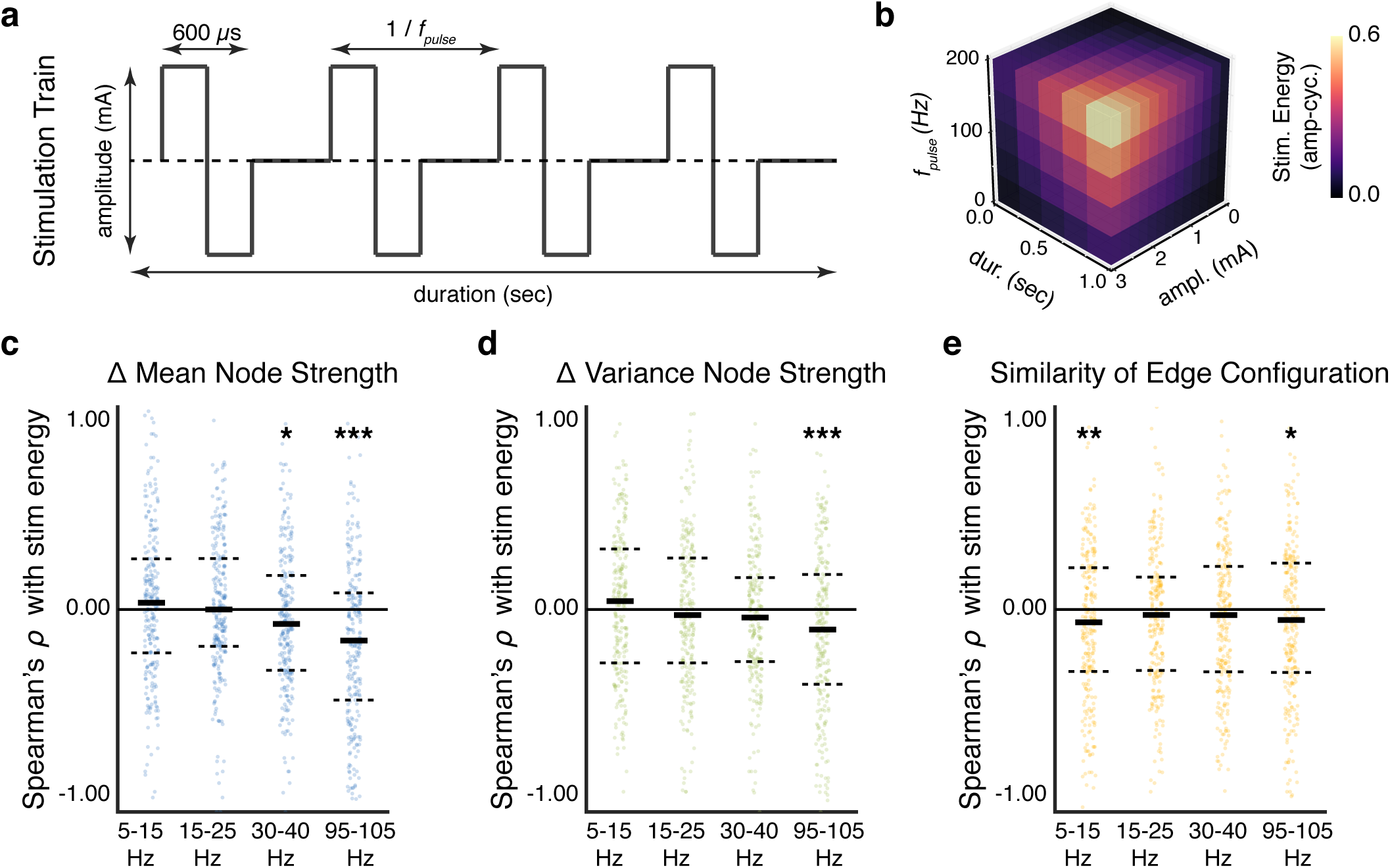
Dose-dependent response of network reconfiguration to stimulation. (**a**) To examine the effect of stimulation energy on network reconfiguration, we vary the amplitude, pulse frequency, and duration of the square-wave input. (**b**) We quantify the total input energy delivered during a stimulation trial as the product between the amplitude, pulse frequency, and duration. Here, we show the three-dimensional plane of input parameters that contribute to the overall stimulation energy. (**c**) Distribution of correlations between the stimulation energy and the change in mean of node strengths. Correlations are significantly negative in the low gamma band (*p* < 0.05, corrected) and in the high gamma band (*p* < 0.001, corrected). (**d**) Distribution of correlations between the stimulation energy and the change in variance of node strengths. Correlations are significantly negative in the high gamma band (*p* < 0.001, corrected). Greater stimulation energy decreases node-level interactions in high-frequency networks and leads to a more homogenous distribution of node strengths in the network. (**e**) Distribution of correlations between the stimulation energy and the configuration similarity. Correlations are significantly negative in the alpha/theta band (*p* < 0.01, corrected) and in the high gamma band (*p* < 0.05, corrected). Greater stimulation energy leads to lower configuration similarity (greater edge reconfiguration) in both the low frequency and high frequency networks. For high frequency networks, the extent of edge-level reconfiguration may subserve a finer-scale mechanism for node-level alterations in functional network topology. Each observation is the correlation across epochs within a stimulation session of a single subject. Solid lines represent the median, and dashed lines represent first and third quantiles. \**p* < 0.05, \*\**p* < 0.01, \*\*\**p* < 0.001.

To test this hypothesis, we first compute a within-session Spearman’s ? correlation between stimulation energy and the three measures of functional network reconfiguration (change in mean of node strengths, change in variance of node strengths, and configuration similarity) for the four coherence frequency bands (**Fig. 3c-e**). Using a one sample *t*-test and Bonferroni correction for multiple comparisons, we test whether increasing stimulation energy drives greater node-level changes in the network 100ms after stimulation offset (**Fig. 3c,d**). We find that greater stimulation energy leads to a significant decrease in the mean of node strengths in the low gamma band (*t*(247) =− 2.5, *p* = 0.048) and in the high gamma band (*t*(247) =− 6.3, *p* = 4.4 10^−9^), and to a non-significant change in the alpha/theta band (*t*(247) = 1.5, *p* = 0.56) and in the beta band (*t*(247) = 0.6, *p* = 1.0). We also find that greater stimulation energy leads to a significant decrease in the variance of node strengths in the high gamma band (*t*(247) =− 5.9, *p* = 5.8 × 10^−8^), and to a non-significant change in the alpha/theta band (*t*(247) = 1.0, *p* = 1.0), in the beta band (*t*(247) =− 0.6, *p* = 1.0), and in the low gamma band (*t*(247) =$#x2212; 1.7, *p* = 0.32). We find that these effects indeed persist in the high gamma band at least 600ms after stimulation offset (**Fig. S3c,d**). Our results indicate a robust dependence of high frequency functional reorganization at the scale of network nodes on stimulation strength. Specifically, greater stimulation energy disrupts and decreases cohesive node-level interactions in high frequency bands. We did not observe a similar disruption in node-level architecture in the lower frequency bands.

Logically, we next ask whether stimulation energy similarly alters the edge-level architecture of the network. Using a one sample *t*-test and Bonferroni correction for multiple comparisons, we test whether increasing stimulation energy drives greater configurational change in the network edges 100ms after stimulation offset (**Fig. 3e**). We find that greater stimulation energy leads to a significant decrease in the configuration similarity (greater reconfiguration) in the alpha/theta band (*t*(247) =− 3.2, *p* = 0.005) and in the high gamma band (*t*(247) =− 2.0, *p* = 0.044), and to a non-significant change in the beta band (*t*(247) =− 1.7, *p* = 0.33) and in the low gamma band (*t*(247) =− 1.0, *p* = 1.0). We find that these effects dissipate 600ms after stimulation offset (**Fig. S3e**). Our results indicate that greater stimulation energy drives greater reconfiguration of the functional topology in both low frequency bands and high frequency bands. Critically, we find that stimulation strength only explains immediate edge-level reconfiguration of network topology and does not exhibit a relationship with later stage edge-level reconfiguration.

Combined with our earlier findings, our analysis demonstrates that greater stimulation energy alters high frequency network architecture at the scale of network nodes and network edges. Specifically, we find that the strength of stimulation drives decreased high frequency coherence between network nodes, by presumably redistributing coherent edges across the network and reducing the variance in node strengths. Conversely, less stimulation energy may increase node strengths in high frequency networks by driving less topological reconfiguration of the network edges and simply reinforcing existing functional interactions. In the low frequency, alpha/theta network, stimulation energy did not significantly influence node-level organization but did drive greater reconfiguration of finer-scale edge architecture – suggesting that low-frequency node-level architecture may be hypersensitive to stimulation to the extent that stimulation strength plays a minimal role.

### 2.3. Stimulation of baseline hubs vs. nonhubs has differential effects on the network

We next build upon our analysis of the influence of stimulation parameters on functional network topology by similarly investigating the role of 248 unique stimulation locations over 94 subjects in the functional brain network (83 depth locations and 165 surface locations, **Fig. 4a**; see **Fig. S5** for regional distribution of stimulation location). We specifically ask, “Do functional hubs drive more widespread reconfiguration of the functional network than functionally isolated brain areas?” To answer this question, we measure the node strength as the mean coherence of the stimulation node to all other nodes at baseline. We hypothesize that stimulation of a stronger functional hub will lead to greater dispersion of input energy throughout the network, driving a homogenous network response; stimulation of a weaker functional hub will lead to more targeted dispersion of input energy to a subset of network nodes, driving a heterogenous network response (**Fig. 4b**).

**Figure 4:**
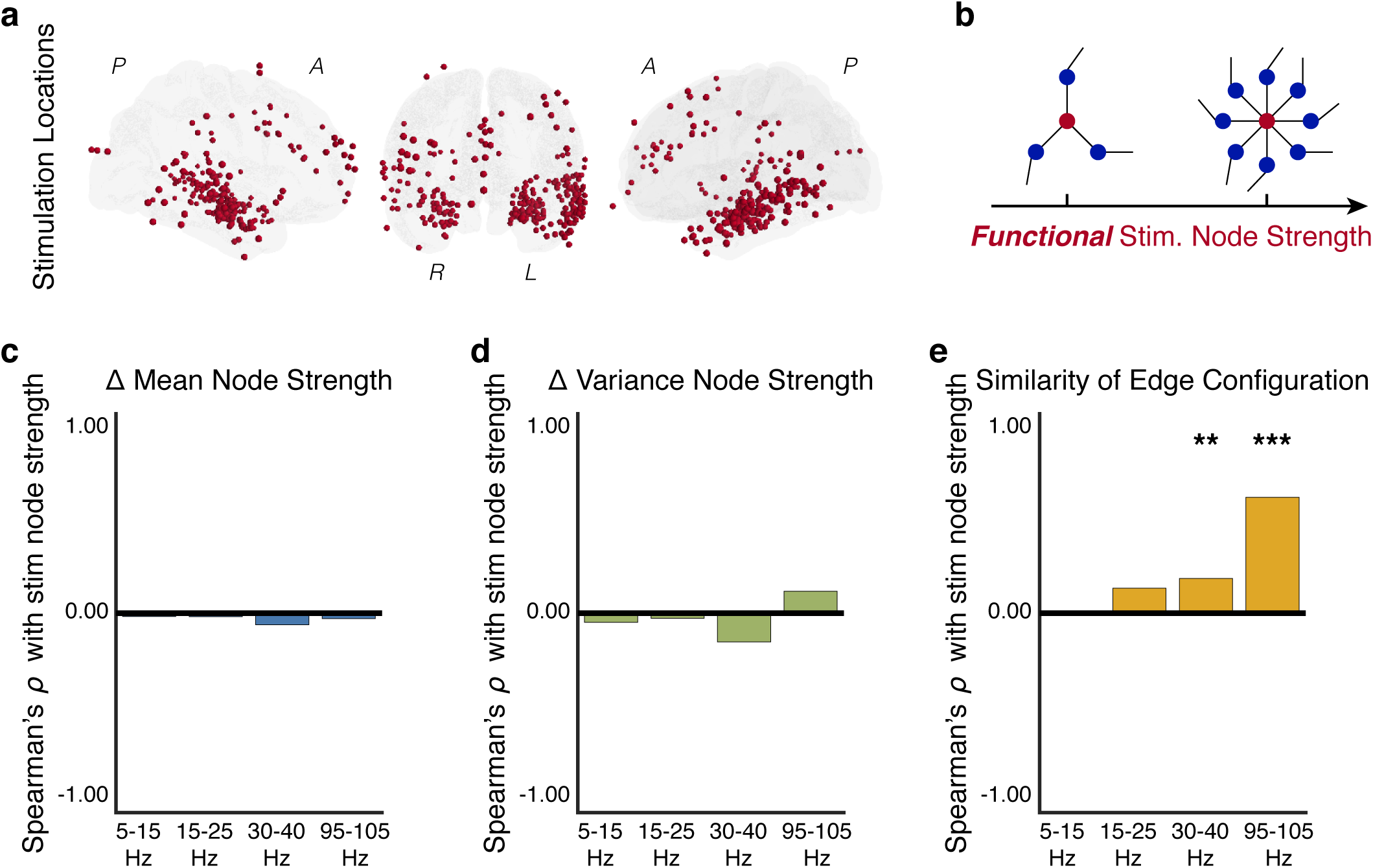
Functional hubs constrain topological response to stimulation. (**a**) Distribution of 248 stimulation locations sampled across 94 subjects. (**b**) To examine the effect of stimulation location on the reconfiguration of functional network topology, we measure the node strength of the stimulation region during the baseline epoch – before any stimulation is delivered – for each coherence frequency band. Intuitively, nodes with low strength (*left*) tend to be functionally isolated and exhibit weak coherence with the other nodes in the network, while nodes with high strength (*right*) tend to be functional hubs and exhibit strong coherence with the other nodes in the network. We expect that stimulation of strong functional hubs will lead to a homogenous change in network topology, and we conversely expect that stimulation of weak functional hubs will lead to a heterogenous change in network topology. (**c**) Correlation between the stimulation node strength and the change in mean of node strengths. We find no significant relationship between the stimulation node strength and the change in mean of node strengths in any frequency band. (**d**) Correlation between the stimulation node strength and the change in variance of node strengths. We find no significant relationship between the stimulation node strength and the change in variance of node strengths. (**e**) Correlation between the stimulation node strength and the configuration similarity. Correlations are significantly positive in the low gamma band (*p* < 0.05, corrected) and in the high gamma band (*p* < 0.001, corrected). Greater stimulation node strength leads to greater configuration similarity (lower edge reconfiguration) in high frequency networks. Correlations are computed over stimulation sessions across subjects. \**p* < 0.05, \*\**p* < 0.01, \*\*\**p* < 0.001.

To test this hypothesis, we first compute a Spearman’s ? correlation between baseline node strength of the stimulation region and the average of each of the three measures of functional network reconfiguration (change in the mean of node strengths, change in the variance of node strengths, and the configuration similarity) for the four coherence frequency bands (**Fig. 4c-e**). Using a Bonferroni correction for multiple comparisons, we test whether greater node strength of the stimulation region drives greater node-level changes in the network 100ms after stimulation offset, over stimulation sessions across subjects (**Fig. 4c,d**). We find that stimulation node strength does not significantly influence the mean of node strengths in the alpha/theta band (ρ(246) = −0.02, *p* = 1.0), in the beta band (ρ(246) = −0.02, *p* = 1.0), in the low gamma band (ρ(246) =− 0.06, *p* = 1.0), or in the high gamma band (ρ(246) =− 0.03, *p* = 1.0). We also find that the stimulation node does not significantly influence the variance of node strengths in the alpha/theta band (ρ(246) =− 0.05, *p* = 1.0), in the beta band (ρ(246) =− 0.03, *p* = 1.0), in the low gamma band (ρ(246) =− 0.14, *p* = 0.09), or in the high gamma band (*p*(246 =− 0.11, *p* = 0.3). We also do not observe these effects after 600ms following stimulation offset (**Fig. S4c,d**). Our results indicate that baseline node strength does not play an influential role in altering large-scale organization of network nodes.

We next ask whether the strength of the stimulation node can differentially drive reconfiguration of edge-level architecture of the network. Using a Bonferroni correction for multiple comparisons, we test whether greater node strength of the stimulation region drives greater configurational change in the network edges 100ms after stimulation offset, over stimulation sessions across subjects (**Fig. 4e**). We find that greater stimulation node strength leads to a significantly greater configuration similarity (lower reconfiguration) in the low gamma band (ρ(246) = 0.17, *p* = 0.03) and in the high gamma band (ρ(246) = 0.58, *p* = 2.2 × 10^−22^), and to a non-significant change in the alpha/theta band (ρ(246) = 0.01, *p* = 1.0) and in the beta band (ρ(246) = 0.13, *p* = 0.2). We find that these effects persist in the low gamma band and in the high gamma band, and that they strengthen in the beta band at least 600ms after stimulation offset (**Fig. S4e**). These results suggest that the functional topology of the stimulation region significantly impacts the pattern of coherent interactions in low and high gamma coherence frequency bands. Specifically, stimulation of weaker functional hubs tends to drive a greater change in the pattern of coherent interactions in low gamma networks and in high gamma networks. Critically, we find that a location-based rule for using stimulation to control the distributed reconfiguration of functional interactions is most robust for high gamma networks thought to reflect activity associated with synaptic input and short-range interactions.

Combined with our earlier findings on the negative relationship between stimulation energy and edge reconfiguration, our findings suggest that stimulation of stronger functional hubs may lead to greater attenuation of the stimulation energy and drive less edge-level reconfiguration than stimulation of weaker functional hubs. Another possible explanation for our findings is that stronger coherent interactions between stimulated hub nodes and the remaining nodes in the network mechanistically constrain the network response to stimulation – which we assess next.

### 2.4. Baseline coherence of stimulation target with other regions constrains future network response

Our findings in the previous section point to an important role of the baseline functional network in constraining the network response to stimulation. Logically, a stimulation node that exhibits stronger coherence with one set of brain regions may be more likely to convey the stimulation input to these brain regions than to another set of brain regions with which it exhibits weaker coherence. We therefore next test the hypothesis that the baseline coherence between the stimulation node and a downstream node predicts the probability that the downstream node will be evoked during a stimulation trial (**Fig. 5a**). In other words, a stronger baseline coherence between the stimulation node and the downstream node may facilitate a greater magnitude change in the node strength of the downstream node.

**Figure 5:**
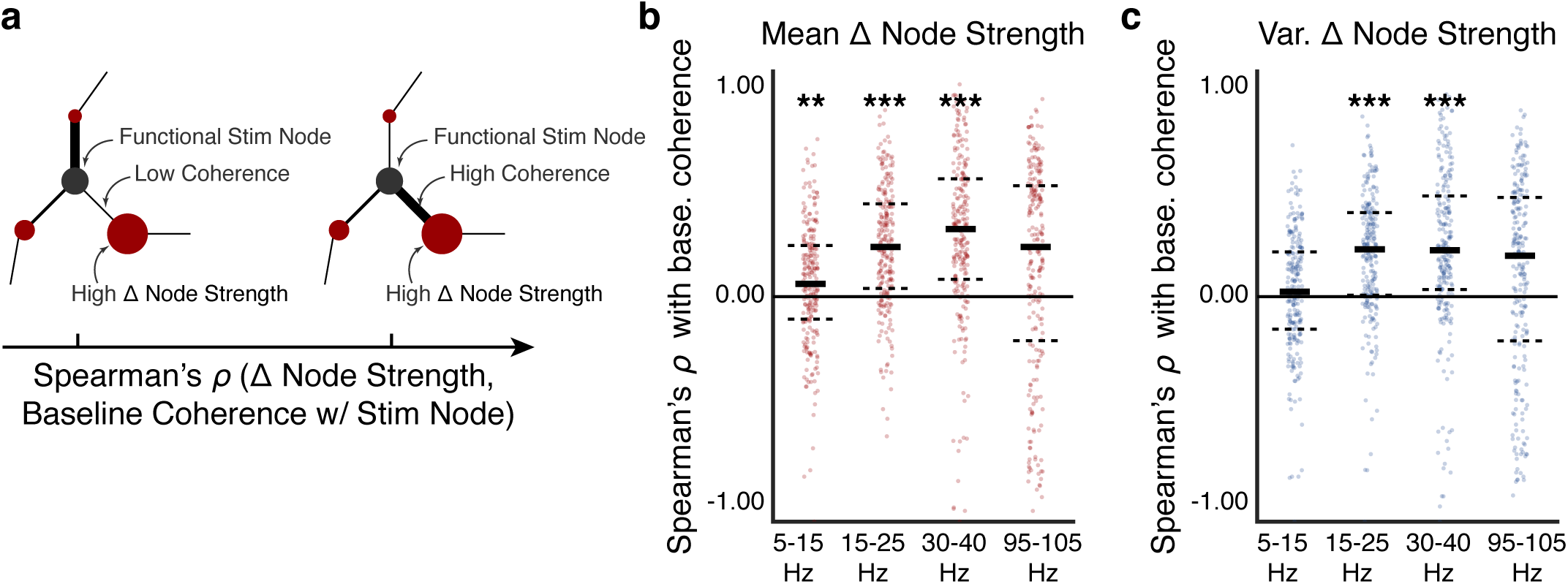
Predicting downstream modulation of regional coherence. (**a**) We hypothesize that the baseline strength of coherent interactions between the stimulation node (red) and network nodes away from the stimulation node (black) predicts the likelihood that these downstream nodes will be evoked due to stimulation. Intuitively, a weak baseline coherence between the stimulation node and a downstream network node is less likely to modulate the mean coherence of the downstream node (*left*), and a strong baseline coherence between the stimulation node and a downstream network node is more likely to modulate the mean coherence of the downstream node (*right*). To test this hypothesis, we first quantify the magnitude change in node strength within each stimulation session. We next compute Spearman’s ? correlation between the mean (and variance) of change in downstream node strength across stimulation trials and the baseline coherence between the stimulation node and the downstream nodes. (**b**) Distribution of correlations between the mean of change in downstream node strength and the baseline coherence between the stimulation node and the downstream nodes, for each of the four coherence frequency bands. We find a significantly positive correlation in the alpha/theta band (*p* < 0.01, corrected), in the beta band (*p* < 0.001, corrected), and in the low gamma band (*p* < 0.001, corrected). These results suggest that baseline functional network topology involving the stimulation node predicts downstream modulation in node strength. (**c**) Distribution of correlations between the variance of change in downstream node strength and the baseline coherence between the stimulation node and the downstream nodes. We find a significantly positive correlation in the beta band (*p* < 0.001, corrected) and in the low gamma band (*p* < 0.001, corrected). These results suggest that baseline functional network topology involving the stimulation node predicts the flexibility with which a downstream node may alter its interactions with other nodes in the network. Each observation is the correlation within a stimulation session of a single subject. Solid lines represent the median, and dashed lines represent the first and third quantiles. \**p* < 0.05, \*\**p* < 0.01, \*\*\**p* < 0.001.

To address this hypothesis, we first compute the within-session mean and variance of the magnitude change in node strength. For each session, we next compute the Spearman’s ? correlation between the set of baseline coherence values between the stimulation node and the downstream nodes and the mean of the magnitude changes in node strength of each downstream node (**Fig. 5b**). Intuitively, positive correlation values imply that stronger baseline coherence between the stimulation node and the downstream nodes predicts greater magnitude change in node strength across stimulation epochs. Using a Wilcoxon rank-sum test and Bonferroni correction for multiple comparisons, we find a significant positive correlation values in the alpha/theta band (*Z*(247) = 4986, *p* = 0.003), in the beta band (*Z*(247) = 2092, *p* = 5.4 × 10^−15^), in the low gamma band (*Z*(247) = 2355, *p* = 1.4 × 10^−13^) and we find a non-significant positive trend in the high gamma band (*Z*(247) = 5928, *p* = 0.19). We also assess the Spearman’s ? correlation between the set of baseline coherence values between the stimulation node and the downstream nodes and the variance of the magnitude changes in node strength of each downstream node (**Fig. 5b**). We find a significantly positive correlation in the beta band (*Z*(247) = 2721, *p* = 9.9 × 0^−11^) and in the low gamma band (*Z*(247) = 2997, *p* = 2.0 × 10^−10^), and we find a non-significant positive trend in the alpha/theta band (*Z*(247) = 5993, *p* = 0.24) and in the high gamma band (*Z*(247) = 6340, *p* = 0.72). We find that these effects persist in the alpha/theta band, in the beta band, and in the low gamma band and strengthen in the high gamma band at least 600ms after stimulation offset (**Fig. S6b,c**).

Our findings are consistent with the hypothesis that the baseline functional network topology involving the stimulation node may be used to predict the downstream network regions that are most influenced by stimulation. We also find that this predi-ctive capacity is dependent on the frequency band of the coherent measurement: the likelihood of modulating the coherent interactions of a downstream node is less predictable for higher frequencies. This finding suggests that the direct coherence between a stimulation node and a downstream node may be more influential in conveying stimulation input to the downstream node for lower coherence frequency bands. Our analysis provides critical insight into mechanisms of node-level flexibility, or the ability for a network region to dynamically alter its level of interaction with other regions in the network. Specifically, stronger baseline coherence between the stimulation node and a downstream node tends to predict greater variability with which the downstream node changes its level of interaction with the rest of the network within a stimulation session. Such a rule can guide more principled targeting of network structures to amenably drive flexible reconfiguration of the functional network.

### 2.5. Unifying stimulation and functional reconfiguration with network control theory

We lastly seek to integrate our observations on stimulation-driven reconfiguration of functional brain networks with first principles theory. Network control theory provides a mathematical framework to model changes in the state of a complex system under a set of constraints prescribed by the structure of that system [77, 98]. For brain networks, network control theory offers an opportunity to model the logical progression of a stimulus input into an anatomically-defined structural brain network, the traversal of that input through the network, the resulting change in inter-regional communication, and an accompanying shift in the dynamical state of the brain that accommodates a change in behavior (**Fig. 6a**) [42, 41, 72, 78, 59]. The structural topology of the network may confer important control properties to a complex system such as *modal controllability*, which enables a system to move from its current dynamical state to more difficult-to-reach dynamical states through an efficient expenditure of energy resources [42, 77, 4]. Recent theoretical inquiry into the relationship between brain structure and function during stimulation posited that stimulation of the structural brain network’s modal controllers may drive a heterogenous change in functional architecture [72]. However, experimental evidence linking the network control theoretic model to brain stimulation and its influence on functional architecture and dynamical brain state via the structural brain network is lacking.

**Figure 6:**
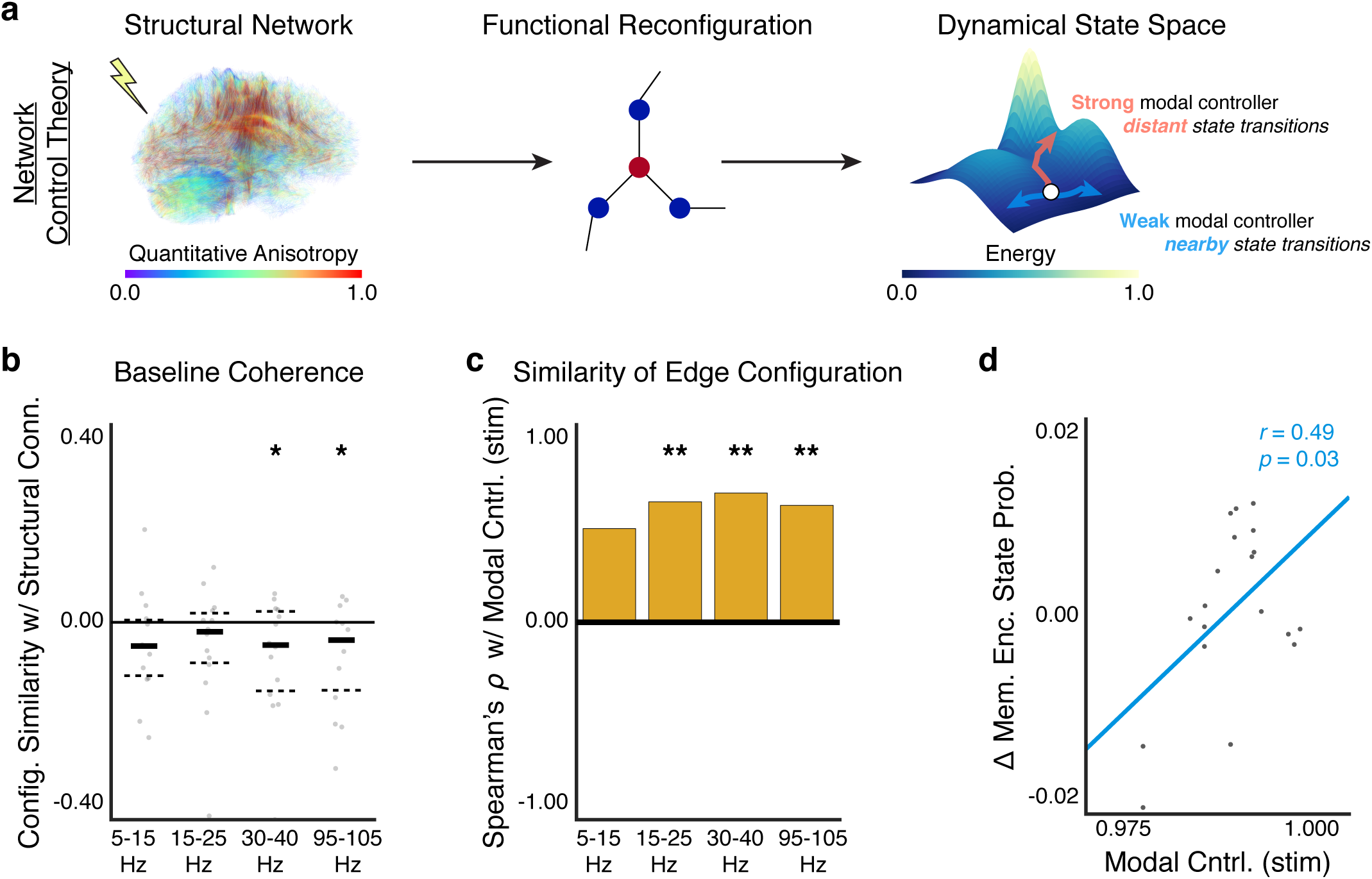
Using neurostimulation to bridge structure, function, and behavior. (**a**) Network control theory can model dynamical state changes due to external input and structural constraints on the system. Consider a stimulation input to the structural brain network (*left*). This input would evoke a functional response constrained by the architecture of the structural network (*middle*) and shift the brain from one state to another state (*right*). Previous studies posit that structural topology of the input region – modal controllability – plays a critical role in the energy required to move between different dynamical states [42, 77]. Specifically, stronger modal controllers may lead to more distant transitions across an energy landscape than weaker modal controllers (*right*). Here, we demonstrate a critical link between stimulation of the structural brain network, the evoked functional network response, and changes in dynamical state associated with behavior. Briefly, we use a previously published biomarker of brain state based on a logistic regression-based classifier of neural activity associated with positive memory encoding [35]. (**b**) Distribution of the configuration similarity between structural connectivity and baseline functional connectivity in four coherence frequency bands. Each observation is a single subject. Solid lines represent the median, and dashed lines represent the first and third quantiles. We find negative trends in the configuration similarity between structural connectivity and baseline functional connectivity in the low gamma band (*p* < 0.05, uncorrected) and in the high gamma band (*p* < 0.05, uncorrected). (**c**) Correlation between the modal controllability of the stimulated brain region and the average functional configuration similarity across stimulation sessions. We find a significant positive correlation in the beta band (*p* < 0.01, corrected), in the low gamma band (*p* < 0.01, corrected), and in the high gamma band (*p* < 0.01, corrected). This result implies that stimulation of modal controllers leads to less network-wide reconfiguration of functional interactions. (**d**) We find a significant positive correlation between the average change in classifier likelihood of positive memory encoding state during stimulation trials and the modal controllability of stimulated nodes based on the structural brain network (*p* < 0.05). This result implies that stimulation of structural brain regions that are more capable of pushing the brain to difficult-to-reach dynamical states is associated with an increased likelihood of reaching a positive memory encoding state after stimulation. \**p* < 0.05, \*\**p* < 0.01, \*\*\**p* < 0.001.

In this study, we have so far shown that stimulation drives a rich and complex functional network response that is dependent on the stimulation input energy and the stimulation input location. While the input energy drives the magnitude of functional reconfiguration, the baseline coherence of the input location constrains the spatial specificity of the functional reconfiguration. Yet, how does baseline network topology in the vicinity of the stimulation region relate to the modal control strategy put forth by structural control theory? To answer this question, we construct structural brain networks by applying deterministic tractography to diffusion tensor imaging data that are parcellated into 1015 anatomically-defined, cortical and subcortical regions of interest (ROI) [28] in a subset of 14 participants (see *Methods*; the other participants did not have diffusion tensor imaging data). We next assign the intracranial ECoG sensors for a subject to their nearest anatomical ROI based on the shortest spatial distance between a sensor and the ROI centroids. We then measure the structural connectivity between anatomical ROIs and the modal controllability of each anatomical ROI in the structural network, and we assign these values to the intracranial sensors based on their proximity to the nearest ROI.

To establish a clear relation between structural connectivity and baseline functional connectivity (measured by coherence), we calculate the configuration similarity between the pattern of structural connections and the pattern of baseline coherent interactions between intracranial sensors, for each coherence frequency band and each subject (**Fig. 6b**). Using a one sample *t*-test, we test the null hypothesis that the configuration similarity is equalbaseline coherent interactions to zero. We observe non-significant, negative trends between structural connectivity and baseline coherence in the alpha/theta band (*t*(13) = −1.7, *p* = 0.11, uncorrected), in the beta band (*t*(13) =− 1.5, *p* = 0.16, uncorrected), in the low gamma band (*t*(13) =− 2.2, *p* = 0.04), and in the high gamma band (*t*(13) =− 2.4, *p* = 0.03). Our results imply that stronger structural connections are generally associated with weaker coherent interactions, most saliently in higher frequency bands that typically reflect local, bottom-up processing associated with synaptic input.

Based on the result that structural connectivity weakly constrains baseline coherent interactions between brain regions, we ask whether a control strategy motivated by modal controllability of the structural brain network predicts stimulation-driven reconfiguration of the functional network topology. To answer this question, we first calculate the modal controllability for the brain region in each stimulation session across subjects. We next compute the Spearman’s ? correlation between modal controllability of the stimulated brain region and the average, within-session configuration similarity. Using Bonferroni correction for multiple comparisons, we find a significant positive correlation between modal controllability and configuration similarity in the beta band (ρ(24) = 61, *p* = 0.003), in the low gamma band (ρ(24) = 0.65, *p* = 0.001), and in the high gamma band (ρ(24) = 0.59, *p* = 0.005), and we find a nonsignificant positive trend in the alpha/theta band (ρ(24) = 0.47, *p* = 0.05). These effects persist at least 600ms after stimulation offset (**Fig. S7a**). These results imply that stimulation of stronger modal controllers drives lower reconfiguration of the coherent interactions in the functional network. To contextualize these findings, we draw upon the well-established positive relationship between modal controllability and structurally isolated brain regions [42]. By targetting strong modal controllers in the structural brain network, stimulation modulates structurally isolated brain areas and drives lower functional reconfiguration than stimulation of weak modal controllers in structurally connected brain areas.

Notably, our results demonstrate a complex and frequency-specific link between structural topology and functional topology – weak structural connections tend to span between brain areas with stronger, high frequency functional interactions and stimulation of modal controllers in structurally isolated brain regions tends to limit the extent of functional network reconfiguration. Combined with earlier findings, we put forth a putative sequence of physiological events associated with modal control in which stimulation of strong modal controllers activates strong, local functional hubs that drive less functional reconfiguration of network-wide edges (**Fig. 4e**) and greater change in the node strengths of functionally connected, downstream brain regions (**Fig. 5b**). In contrast, stimulation of weak modal controllers activates weaker, functionally isolated regions that drive distributed functional reconfiguration of network-wide edges (**Fig. 4e**). While these findings establish a link between network control and functional reconfiguration, they do not establish a link between network control and changes in dynamical brain state.

To experimentally examine the relationship between the modal controllability of the stimulation region and the shift in dynamical brain state following stimulation, we leverage a previously documented and validated binary classifier of neural activity into good and poor episodic memory encoding states [35]. Specifically, we first train the classifier to discriminate between successful and unsuccessful word recall trials during a delayed free recall task using features based on spectral power of ECoG activity. We next evaluate the classifier on ECoG activity during the pre-stim epoch and the post-stim epoch of each stimulation trial and compute the change in probability of being in a good memory encoding state. We calculate the correlation between the modal controllability of the stimulation region and the average, within-session change in probability of being in a good memory state over stimulation trials (**Fig. 6d**). We find a significant positive correlation (Pearson’s *r*(17) = 0.49, *p* = 0.03). These findings imply that the push towards better memory encoding states is associated with stimulation of strong modal controllers that are theoretically positioned to push the brain to more energetically unfavorable and distant brain states.

Collectively, our study reveals a link between brain structure and brain function that is grounded in network control theory. Using the network control framework, we uncover the important role of stimulation on reconfiguration of functional architecture that accounts for anatomical constraints on network dynamics via the topology of structural network connectivity. Critically, we find that brain networks may use a modal control strategy during transitions between difficult-to-reach dynamical states, which is associated with a reconfiguration in the localized coherence of individual network nodes to the broader functional brain network.

## 3. Discussion

Here, we addressed the hypothesis that direct stimulation of cortical and subcortical structures alters the architecture of functional brain networks and shifts the dynamical brain state in accord with control strategies identified by applying tools from network control theory to electrophysiological and structural brain imaging data. In human epilepsy patients, we measured coherent patterns of ECoG activity thought to underlie coordinated functional interactions and we mapped how these interactions vary with neurostimulation parameters. We observed that stimulation drives two modes of functional reconfiguration: the first mode involves distributed changes in the pattern of functional interactions across the network, and the second mode involves preferentially localized changes in the functional interactions associated with select brain regions. Notably, the mode of reconfiguration may be strategically selected based on the strength and location of stimulation. When we stimulated brain regions with weak structural connections to the rest of the network, we tended to invoke a modal control strategy marked by a modulation of the functional hubness of downstream brain regions and a large change in dynamical brain state.

### 3.1. Predictors of functional reconfiguration: Implications for brain network control

The field of network neuroscience has long sought to understand how the rigid and interconnected anatomy of the structural brain network shapes interactions amongst functionally-specialized brain areas, which may change from moment to moment and drive cognition and behavior [70, 44, 43, 76]. Tools from network control theory [65, 77, 89] have enabled researchers to identify *controllability* rules that prescribe how the dynamical state of a neural system can change based solely on the structural topology of the system [15, 42, 59, 41, 90, 98]. Structural controllability rules account for network interactions that *can* occur, but they do not account for finer-scale functional constraints that dictate whether these interactions *will* occur at a point in time [65, 89]. Previous studies have incorporated these functional constraints into the study of network control by using neurophysiologically-inspired dynamical mean-field models [50, 91, 72], which require estimation of biologically plausible parameters.

In contrast to these studies, we used a data-driven, perturbative approach for inferring rules of functional network reconfiguration. By focally stimulating brain tissue at the millimeter-scale, we mapped changes in the statistical interdependencies between brain regions. We note that there is a subtle distinction in the statistical methods used here to measure bidrectional, synchronized interactions and statistical methods used elsewhere to measure directed, effective functional interactions [39, 64, 14]. We found that the observed change in statistical interdependency can be predicted by baseline levels measured before any stimulation is delivered. That is, a brain region that exhibits a strong, spontaneous functional interaction with the stimulated brain region is more likely to be modulated during stimulation than a brain region with a weak, spontaneous functional interaction with the stimulated brain region. In the long standing debate regarding the validity of functional network models to explain causal dynamics [51], these data provide compelling evidence in favor of a mechanism in which the functional network may causally convey the influence of stimulation on one brain region to other strongly interacting brain regions. Our findings implicate a strategy for the *functional control* of brain networks in which (i) stimulation of functionally isolated brain regions leads to spatially focal and strong downstream functional reconfiguration, and (ii) stimulation of functionally hub-like brain regions leads to spatially diffuse and weak downstream functional reconfiguration.

To relate rules for functional control to theoretical predictions from structural controllability, we bridged electrophysiological data with structural brain imaging data. Confirming results from prior studies on controllability in healthy subjects [42, 90], in epilepsy subjects we found that brain regions with structurally weak connections tend to be strong modal controllers, which facilitate more difficult transitions in brain state.

Stimulations of structurally weak modal control points lead to strong, local change of functional architecture in a mean-field model of neuron population dynamics [72]. In contrast, here we demonstrated that stimulation of modal control points leads to widespread, weak change in functional architecture due to the presence of strong functional hubs in the stimulated brain regions. We identified two potential explanations for the distinction between the model simulation and our empirical observations. First, the *in silico* model assumes identical biophysical parameters across different neuronal ensembles distributed across the brain network, which may limit the reproducibility of detailed spatial and temporal dynamics that would otherwise be expressed *in vivo* and measured by intracranial ECoG sensors. Second, the *in silico* model accounts for structural connectivity between neuronal ensembles using diffusion imaging, which measures white-matter fiber pathways spanning long distances but does not capture gray-matter pathways spanning short distances [92] and synaptic microarchitecture responsible for plasticity over varying time scales [86]. When combined with additional data demonstrating a moderate relationship between white-matter connectivity and correlated ECoG dynamics [16], our findings suggest that non white-matter structural connectivity and other physiological factors may contribute to the reconfiguration of functional network architecture.

### 3.2. Physiological interpretations of altered functional topology

Neuronal synchronization is purported to play a crucial role in facilitating interareal communication between ensembles of neurons [29, 27, 82, 38, 18]. Fries (2015) [38] proposed that rhythmic oscillations in the local field potential give rise to states of excitability depending on the temporal position during an oscillatory cycle – two different ensembles of neurons are able to reliably transfer information between one another when they are mutually excitable, or exhibit oscillations that are inphase. Equally important to communication is the frequency of the oscillation – higher frequency bands (γ) are thought to facilitate communication of bottom-up input over short distances and lower frequency bands (θ, α, β) are thought to facilitate communication of top-down processes over long distances [38, 60].

Notably, we found that stimulation parameters may be tuned to selectively modulate different regions and spatial extents of the functional network. Stimulation energy tends to provide greater control over reconfiguration of lower frequency networks, and stimulation location tends to provide greater control over reconfiguration of higher frequency networks. Specifically, we observed that the strength of the stimulation input has a greater effect on functional reconfiguration in lower frequency bands than in higher frequency bands, suggesting that stimulation parameters such as amplitude, pulse frequency, and duration may play an important role in the modulation of longrange, top-down functional interactions. We speculate that a stronger stimulation input may be more likely to penetrate wider spatial extent of cortex and heterogenously modulate network excitability at low frequencies [38]. We also observed that the location of the stimulation input has a greater effect on functional reconfiguration in higher frequency bands than in lower frequency bands, suggesting that the hubness of the stimulated brain region may play an import role in the modulation of short-range, bottom-up functional interactions. Intuitively, we expect that brain regions involved in bottom-up communication associated with broadly conveying sensory input to higher order cortices might also be more sensitive to modulations via stimulation than brain regions involved in top-down communication.

### 3.3. Methodological considerations and future work

We chose to perform a network analysis of intracranial data during neurostimulation, rather than a univariate analysis of individual activations. Our decision enabled us to examine the influence of neurostimulation on the distributed and interconnected physiology of the human brain, which is in line with previous *in silico* modelling work [72]. A wealth of important studies have teased apart the effective connectivity between brain regions using neurostimulation to generate cortico-cortico evoked potentials (CCEPs) [54, 55, 68]. In a departure from these studies on effective connectivity, we test the novel hypothesis that neurostimulation can predictively perturb interareal statistical dependencies underlying distributed brain function. Our study is motivated by recent hypotheses on the role of coherent synchronization in neuronal communication. With the maturation of network analysis tools that enable simultaneous tracking of dynamic network architecture and dynamic activity [74], we envision future studies where we investigate the potential for neurostimulation to selectively modulate brain activity or perturb functional network architecture.

We also chose to examine static changes in the functional network architecture aggregated over many repeated trials of neurostimulation. This approach gave us the ability to examine the statistical robustness of network reconfiguration – and to address questions such as “Are some brain regions more likely to change their functional interactions than other brain regions?” And “How are these brain regions associated with the stimulated brain region?” In future studies, we aim to understand how stimulation influences the functional brain network from one moment in time to the next, as a function of brain state. For instance, does stimulation during the beginning of a coherent oscillatory cycle influence network architecture differently than stimulation during the middle of a coherent oscillatory cycle? This temporal mapping could inform control strategies to steer functional brain network reconfiguration in real-time.

Approaches for recording and interrogating intracranial electrophysiology are inherently limited by spatial coverage of the ECoG sensors, which is determined during the management of a patient’s epilepsy. This sampling bias leads to a varied representation of the functional brain network between individual patients. While it is not yet possible to record from the entirety of the human brain using ECoG, we mitigated this shortcoming by taking key steps in our analysis. First, we used a statistically robust approach for characterizing the network-impact of stimulation in individual patients. We computed separate measures of topology for each patient’s functional brain network, which enabled us to account for individual variability in sensor placement and physiological state. Our data demonstrated a set of functional rules for network reconfiguration that fundamentally depend on topological characteristics of the stimulated brain area that can vary within and between patients. Second, we used a large dataset consisting of ninety four epilepsy patients, allowing us to account for a range of individual variability in functional brain network architecture that is often not possible in studies of human electrophysiology.

## 4. Conclusions

Here we mapped, for the first time, the impact of targeted neurostimulation on distributed functional architecture in the human brain. We demonstrated that network physiology can be predictably altered based on control theoretic rules that account for structural and functional organization of the brain network. Our results provide a causal, quantified description of the influence of structure and function on dynamical brain state. Our findings have significant translational implications in strategizing stimulation-based therapy based on a combination of behavioral biomarkers and neurophysiology.

## 5. Acknowledgments

ANK and DSB acknowledge support from the John D. and Catherine T. MacArthur Foundation, the Alfred P. Sloan Foundation, the Army Research Laboratory and the Army Research Office through contract numbers W911NF-10-2-0022 and W911NF-14-1-0679, the National Institute of Health (2-R01-DC-009209-11, 1R01HD086888-01, R01-MH107235, R01-MH107703, and R21-M MH-106799), the Office of Naval Research, and the National Science Foundation (BCS1441502, CAREER PHY-1554488, and BCS-1631550). We thank Blackrock Microsystems for providing neural recording and stimulation equipment. This work was supported by the DARPA Restoring Active Memory (RAM) program (Cooperative Agreement N66001-14-2-4032). The views, opinions, and/or findings contained in this material are those of the authors and should not be interpreted as representing the official views or policies of the Department of Defense, the U.S. Government, or any of the funding agencies.

## 6. Methods

### 6.1. Study cohort

Ninety four patients undergoing intracranial EEG monitoring as part of clinical treatment for drug-resistant epilepsy were included in this study. Data were collected as part of a multicenter project designed to assess the effects of electrical stimulation on memory-related brain function. Data analyzed in this study were collected at the following centers: Thomas Jefferson University Hospital (*N* = 23), University of Texas Southwestern (*N* = 23), Mayo Clinic (*N* = 17), National Institutes of Health (*N* = 11), Dartmouth-Hitchcock Medical Center (*N* = 9), Hospital of the University of Pennsylvania (*N* = 6), Columbia University Medical Center (*N* = 4), Emory University Hospital (*N* = 1). The research protocol was approved by the institutional review board (IRB) at each hospital and informed consent was obtained from each participant.

### 6.2. Anatomical Localization of Intracranial Electrodes

Patients undergoing surgical treatment for medically-refractory epilepsy believed to be of neocortical origin underwent implantation of intracranial electrodes to localize the seizure onset zone. These procedures were applied after presurgical evaluation with scalp EEG recording of ictal epochs, MRI, PET and neuropsychological testing suggested that focal cortical resection may be a therapeutic option. Patients were then deemed candidates for implantation of intracranial electrodes to better define epileptic networks. Electrode configurations spanned the surface of the cortex (linear and two-dimensional arrays, each sensor is 2.3 mm diameter spaced 10 mm apart) and subcortical depth (each sensor is 1.5-10 mm apart). All electrode configurations were planned by a multidisciplinary team of neurologists and neurosurgeons at each of the eight medical centers.

Electrodes were anatomically localized using separate processing pipelines for surface and depth electrodes. To localize depth electrodes we first labeled hippocampal subfields and medial temporal lobe cortices in a pre-implant, 2 mm thick, coronal T2-weighted MRI using the automatic segmentation of hippocampal subfields (ASHS) multi-atlas segmentation method [102]. We additionally used whole brain segmentation to localize depth electrodes not in medial temporal lobe cortices. We next co-registered a post-implant CT with the pre-implant MRI using Advanced Normalization Tools (ANTs) [5]. Electrodes visible in the CT were then localized within sub-regions of the medial temporal lobe by a pair of neuroradiologists with expertise in medial temporal lobe anatomy. The neuroradiologists performed quality checks on the output of the ASHS/ANTs pipeline. To localize subdural electrodes, we first extracted the cortical surface from a pre-implant, volumetric, T1-weighted MRI using Freesurfer [36]. We next co-registered and localized subdural electrodes to cortical regions using an energy minimization algorithm [34]. For patient imaging in which automatic localization failed, the neuroradiologists performed manual localization of the electrodes.

### 6.3. Electrophysiological Data Acquisition and Stimulation Mapping Protocol

The electrocorticogram (ECoG) was recorded and digitized at 500 Hz, 512 Hz, 1000 Hz, 1024 Hz, or 2000 Hz depending on clinical considerations at each medical center. Signals were recorded using a referential montage with the reference electrode, chosen by the clinical team, distant to the site of seizure onset.

To study the response of the electrocorticogram to neurostimulation, we used a mapping procedure in which stimulation was delivered to cortical and subcortical brain regions – patients were not instructed to engage in any other task before or during stimulation. Prior to the start of each mapping session, we selected a pair of adjacent electrodes for stimulation by prioritizing electrodes in brain regions thought to be associated with memory function. For each mapping session, we selected a new stimulation site and patients underwent one or several mapping sessions depending on their availability for testing and the monitoring needs of the clinicians. Prior to the start of a mapping session, we recorded thirty seconds of ECoG activity as a *baseline* epoch. During a mapping session, we performed several stimulation trials in which a single trial consisted of the following epochs: (i) a half-second pre-stimulation epoch, (ii) a stimulation epoch with variable duration, two consecutive and non-overlapping half-second post-stimulation epochs, and an inter-stimulation epoch with variable duration. During each stimulation trial, we delivered stimulation using charge-balanced, biphasic, rectangular pulses with a pulse width of 300 ms and the following parameters uniformly selected from discrete distributions: pulse frequency (10, 25, 50, 100, 200 Hz), pulse amplitude (maximum safe amplitude minus 0, 0.5, 1 mA; range of 0.125–3.0 mA across subjects), stimulation duration (250, 500, 1000 ms), and inter-stimulation interval (2750–3250 ms). These stimulation parameter ranges were chosen to be well below the accepted safety limits for charge density [83] and ECoG was continuously monitored for afterdischarges by a trained neurologist.

To eliminate confounding effects of stimulation on signal quality and saturation, we disregarded ECoG data collected during the stimulation epoch and the 100 msec following stimulation offset. We also employ a conservative electrode screening procedure, in which we discard non-stimulated channels that exhibit evidence of stimulation-related artifact. Specifically, before re-referencing to a common average reference, we use a paired *t*-test to compare the distribution of mean signal amplitude during the pre-stimulation epoch to the distribution of mean signal amplitude during the post-stimulation epoch, for each electrode across stimulation trials. Using a Bonferroni uncorrected *p*-value threshold of 0.05, we discard electrodes that exhibit significantly elevated raw, mean signal amplitude during each stimulation session.

We analyzed ECoG data collected during the baseline, prestimulation, and post-stimulation epochs. The post-stimulation epoch following 100ms of a buffer period, was split into two consecutive and non-overlapping segments, 0.5 seconds in duration to assess delayed effects of stimulation – we refer to the first segment as the 100ms response and the second segment as the 600ms response. We compared stimulation-related change in neural activity to spontaneous change in neural activity by dividing the baseline epoch into segments equal in duration as the pre-stimulation segment and the post-stimulation segments. To control for the effect of time, we re-sampled segments with time-spacing equal to the length of time between the end of the pre-stimulation epoch and the start of the post-stimulation epoch. To account for the impact of stimulation current on the recording properties of the intracranial sensors, we removed the stimulated electrodes from all analyses of the pre-stimulation and post-simulation epochs – we retained the stimulated electrodes for all analyses of the baseline epoch.

### 6.4. Constructing Frequency-Based Functional Brain Networks

ECoG signals were divided into 0.5s, non-overlapping, time windows – the baseline epoch consisted of sixty time windows spanning thirty seconds, the pre-stimulation epoch consisted of one time window per stimulation trial, and the post-stimulation epoch consisted of two time windows per stimulation trial (100-600ms post-stimulation and 600ms-1100ms post-stimulation). We applied a common average reference to the artifact-free ECoG signal before constructing functional networks [93, 62, 63, 24, 57].

To measure functional interactions between ECoG signals in each time window, we computed spectral coherence, which is a measure of correlation between the power spectra of two signals within a frequency range. Prior studies have shown that coherence is largely independent of the shape of the power spectrum in ECoG signals [23, 22, 93], and underlies different forms of synchronous interactions between neural populations [60]. We constructed functional networks in each time-window using multitaper coherence estimation, which defines a graph edge between electrode pairs (graph nodes) as the power spectral similarity of signal activity over a specific frequency band. We applied the *mtspec* Python implementation [79] of multitaper coherence estimation with time-bandwidth product of five and eight tapers in accord with related studies [63]. This procedure resulted in a symmetric adjacency matrix **A**(*t*, *f*) with size *N × N*, where *N* is the number of network nodes, or electrode sensors, *t* is the time window, and *f* is the frequency band. In this study, we examined network activity in the following four frequency bands: α/θ; (5–15 Hz), β (15–25 Hz), low-γ (30–40 Hz), high-γ (95–105 Hz). These frequency ranges cover traditional oscillatory classes and have been previously examined for their network topology [63, 57].

An alternate representation of the symmetric, square adjacency matrix **A**(*t*, *f*) is a configuration vector Â(*t*, *f*), which tabulates all *N × N* pairwise interactions. Due to symmetry of the adjacency matrix, we unravel the upper triangle of **A**, resulting in the weights of 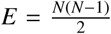 functional interactions. Thus,Â(*t*, *f*) is a vector of size *E*.

### 6.5. Metrics of Functional Network Topology

In this study, we investigated the effect of neurostimulation on functional network architecture at the scale of network nodes and at the scale of network edges. At the node scale, we first quantified the change in the node strength – a measure of functional “hubness” – of individual network nodes. Specifically, we computed the node strength as 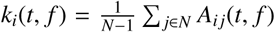, where *k* is the strength of node *i* and *A*_*ij*_ is ^−^the edge weight between nodes *i* and *j*. Based on the time-dependent set of node strengths in the network, we computed the change in the mean of node strengths between time windows and the change in the variance of node strengths between time windows. To assess the magnitude of change in node strength for a node between time windows *t*_*n*_ and *t*_*m*_, we calculated 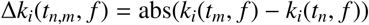.

At the edge scale, we quantified the amount of change in the configurational pattern of the network edges, or coherences, as described previously in [58]. Specifically, we computed the configuration similarity between configuration vectors Â(*t*_*n*_, *f*) and Â(*t*_*m*_, *f*), where *t*_*n*_ and *t*_*m*_ are two different time windows, using the Pearson correlation test statistic. Two vectors with a Pearson correlation value closer to 0 are more dissimilar in their configurational pattern of network edges than two vectors with a Pearson correlation value closer to 1.

For the stimulation epoch, we computed global and local network metrics between the pre-stimulation time window and the post-stimulation time window of a stimulation trial. For the baseline epoch, we computed global and local network metrics between time windows separated by an equal length of time as the duration of stimulations in the associated stimulation session.

### 6.6. Diffusion Tensor Imaging Acquisition and Preprocessing

We collected diffusion tensor imaging data for a subset of patients from Thomas Jefferson University Hospital (*N* = 11) and Hospital of the University of Pennsylvania (*N* = 3) and validated our analysis of the functional network response to neurostimulation.

All scans at Thomas Jefferson University Hospital were acquired with a 3T Philips Achieva with an 8-channel head coil using an echo-planar diffusion-weighted technique. The diffusion scan was 62-directional with a *b*-value of 3000s/mm^2^ and TE/TR = 98/7251 ms. The matrix size was 96×96 and the slice number was 52. The field of view was 23×230mm^2^ and the slice thickness 2.5mm. Acquisition time was 496 sec per DTI scan.

All scans at the Hospital of the University of Pennsylvania were acquired with a 3T Siemens Tim Trio with a 32channel head coil using an echo-planar diffusion-weighted technique. The diffusion scan was 116-directional with a *b*value of 2000s/mm^2^ and TE/TR = 117/4180 ms. The matrix size was 96× 96 and the slice number was 92. The field of view was 210 × 210mm^2^ and the slice thickness 1.5mm. Acquisition time was 506 sec per DTI scan.

Based on recent evidence that diffusion imaging is highly sensitive to subject movement [101] and to directional eddy currents [49], we processed data using the FMRIB Software Library [48]. We first created individual masks of the patient brain using BET [67]. We next simultaneously corrected for motion effects and eddy current distortions by applying the EDDY correction tool [3] to the diffusion scans and a *b* = 0 image collected at the beginning of the scan.

We next reconstructed orientation density functions (ODFs) of the diffusion imaging in each voxel. Specifically, we used DSI Studio (http://www.dsi-studio.labsolver.org) and generalized *q*-sampling imaging (GQI) [100] to compute the quantitative anisotropy (QA) [99] in each voxel. To conduct fiber tractography on the reconstructed diffusion images, we used DSI Studio to generate 1,000,000 streamlines with a maximum turning angle of 35° [6] and a maximum length of 500mm [32]. We next defined the structural brain network using the stream-lines linking *N* = 1015 large-scale cortical and subcortical regions extracted from the Lausanne atlas included in the Connectome Mapping Toolkit [28], consistent with previous work [7, 6, 44, 43, 42, 41, 72]. We summarized these measurements in a symmetric and weighted structural adjacency matrix **S** whose entries *S* _*ij*_ reflect the structural connectivity (quantitative anisotropy) between region *i* and region *j*.

We localized electrodes in native subject T1-weighted MRI space to the Lausanne anatomical space by using ANTs [5] to register the subject’s T1 image to the subject’s diffusion B0 image via affine transformation and also register the subject’s T1 image to MNI space (also native Lausanne space) using a nonlinear warp.

### 6.7. Metrics of Structural Controllability

To study the architectural constraints of the structural brain network with the functional network response to neurostimulation, we adopted a control theoretic approach known as *network controllability*. Briefly, the controllability of a networked system refers to its ability to be driven to specific dynamical states upon external input [53]. Recent research efforts have made substantial progress in the development of quantitative heuristics to characterize different strategies for control [77, 78]. These approaches are now being applied to brain imaging data to understand how structural brain network topology constrains function and behavior [42, 15, 41, 90, 59].

In line with these prior studies, we employ a simplified noisefree linear discrete-time and time-invariant model of network dynamics:

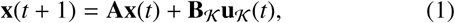

where **x**: ℝ_≥0_ → ℝ^*N*^ describes the state (i.e. voltage, firing rate, BOLD signal) of brain regions over time. Thus, the state vector **x** has length *N*, where *N* is the number of brain regions in the connectome parcellation, and the value of **x**_*i*_ describes the brain activity state of that region. The diagonal elements of the matrix **A** satsify *A*_*ii*_ = 0. Prior to calculating controllability values, we divide **A** by 1 + ξ_0_(**A**), where ξ_0_(**A**) is the largest singular value of **A**. The input matrix 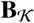 identifies the control point 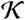 in the brain, where 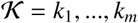 and

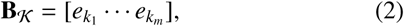

and *e*_i_ denotes the *i*-th canonical vector of dimension *N*. The input 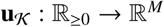 denotes the control strategy.

One control strategy that we investigate in this study is *modal controllability* – the ability of a network region to feasibly control all the dynamical modes of a system [77]. To calculate the modal controllability of an anatomical brain region, we first computed the eigenvector matrix **V** = [*v_i j_*] of the structural network adjacency matrix **S** – intuitively, *v*_*i* *j*_ encodes the ability to control the *j*-th dynamical mode from region *i* [52]. Based on our previous work, we defined 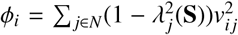 as a scaled measure of the controllability of all *N* dynamical modes λ_1_(**S**), …, λ_*N*_ (**S**) from brain region *i* [77, 42, 72, 90]. Brain regions with high modal controllability are versatile in their ability to control all dynamical modes of the network and brain regions with low modal controllability are specific in their ability to control a subset of dynamical modes of the network.

To provide additional insight into the topological properties of structural control points, we evaluated the structural “hubness” of each brain region by computing the structural node strength as 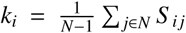 – similar to the calculation for functional node strength specified earlier.

### 6.8. Mapping Intracranial Electrodes to Anatomical Brain Regions

To relate structural controllability to functional network topology of the stimulated electrodes, we first computed metrics of the structural network topology for 463 brain regions defined by the Lausanne anatomical parcellation. The advantage of computing these measures using the anatomical parcellation is the ability to account for whole-brain structural connectivity, including areas that are not directly sampled by the intracranial electrodes. We next assigned intracranial electrodes to the Lausanne brain regions based on a nearest voxel approach. Specifically, we identified the voxel closest to the electrode and assigned the electrode to the brain region containing that voxel. Based on this assignment, we associated values of each structural network metric to the intracranial electrodes.

### 6.9. Detection of Brain States Associated with Memory Encoding

We examined stimulation-driven changes in dynamical brain state using a classifier of neural activity associated with memory encoding processes that was previously validated on data collected during behavioral experimentation with the same patients recruited in this study [35, 61]. Briefly, in these prior studies a logistic regression classifier was trained to discriminate memory encoding-related changes in spectral power in eight logarithmically-spaced frequency bands across intracranial electrodes that are predictive of whether a word was later remembered or forgotten during a free-recall task [35, 61]. In this study, we evaluated the trained memory encoding state classifier on task-free stimulation data of the same patients by measuring spectral power during the pre-stimulation epoch and the post-stimulation epoch, and by computing the change in probability of good memory encoding state for each stimulation trial. We next calculated the average change in probability of good memory encoding state across all stimulation trials of each stimulation mapping session of each patient.

